# Sex-related differences in healthy aging: changes in neuroelectric brain activity reconstructed from resting-state MEG

**DOI:** 10.64898/2026.05.06.723197

**Authors:** Mikhail Ustinin, Anna Boyko, Stanislav Rykunov

## Abstract

Sex-related differences in the aging of the human brain were studied using large array of experimental data. The open archive CamCan was used as a source of data: the magnetic encephalograms, co-registered with magnetic resonance images of the head, were obtained for each of 434 subjects (ages 18-87 years, mean age 54.7 ±18.4): 217 females (ages 18-87 years, mean age 54.5 ±18.4) and 217 males (ages 18-84 years, mean age 54.8 ±18.3). Recordings were split in 10-year age cohorts, each cohort consisted of equal number of men and women to calculate average intersex characteristics correctly. By massively solving the inverse problem, functional tomograms were calculated - the spatial distribution of elementary spectral components. Physiological noise was eliminated by joint analysis of MEG-based functional tomogram and magnetic resonance image for each subject. Then multichannel spectra were transformed into time series of the power of elementary current dipoles. Summary electric powers were calculated in six conventional frequency bands (1-4 Hz – delta; 4-8 Hz – theta; 8-13 Hz – alpha; 13-21 Hz – beta1; 21-30 Hz – beta2; 30-48 Hz – gamma), and sex differences in age-related changes were examined. It was found that in the youngest age cohort (18-29 years) the summary electrical power of the brain for males is 1.5 times greater than such power for females. For adults (30-69 years), male and female powers are approximately equal, while in older cohorts (70-87 years), male total brain power is greater. Age dependencies in various frequency bands are generally different for men and women, excluding higher frequencies 21-48 Hz. Basic conclusion can be made that after intersex averaging total electric power of the human brain is invariant through the lifespan from 18 to 87 years. The proposed method of joint MEG and MRI analysis can be used for further study of the sex-related details of brain sources in their connection with age changes.

## Introduction

Sex and age differences in the human brain are widely studied in various aspects: anatomical, functional, diagnostic, and others. The main method for analyzing the anatomical details of the brain’s structure is magnetic resonance imaging (MRI). It was found that the ratio of total brain volume (TBV) in males and females (TBV_m_/TBV_f_) changes throughout life from 114% in the first age cohort (18–29 years) [Lenroot, 2010] to 110% in the oldest cohorts (44-77 years) [Ritchie 2018]. Based on the results of numerous studies [Ruigrok, 2014], it can be concluded that the male and female brains life trajectories differ anatomically [Bethlehem, 2022, Vucic, 2026] and functionally, operating under different hormonal conditions [Ballard, 2022]. Of interest is the question of age- and sex-related differences in the functioning of the brain as a set of electrical sources. This question is studied using various methods of multichannel encephalography, including approaches such as magnetic encephalography (MEG) [Hoshi, 2020], electroencephalography (EEG) [Han, 2025], and intracranial electroencephalography (icEEG) [Woodhouse, 2025]. Recently, a method of functional tomography based on magnetic encephalography was proposed, based on Fourier transform of the long time series and exploiting mass solution of the inverse problem [Llinás and Ustinin, 2014; Llinás et al., 2015; Ustinin, Boyko, and Rykunov, 2023]. This method presents all registered electrical activity of the head as a large set of elementary current dipoles and was successively applied to the study of age-related human brain changes. It was found that the electrical power of the brain, when averaged across a large mixed population of males and females, remains constant throughout a person’s life, from 18 to 87 years [Ustinin et al, 2025]. The purpose of the present work was to study the sex-related differences in healthy aging changes of the human brain electric power. Changes in the conventional brain rhythms were studied, while electric power was estimated by the inverse problem solution, after the cleaning of the data by joint analysis of the MEG-based functional tomograms and magnetic resonance images.

## Materials and methods

### Data acquisition

In this study we used MEG and MRI data downloaded from CamCAN [Shafto et al., 2014; Henson & Cam-CAN, 2026] data repository. The Cambridge Center for Aging and Neuroscience Research (Cam-CAN) Phase 2 cohort study archive is a large-scale (approximately 700 subjects), multimodal (MRI-magnetic resonance imaging, MEG-magnetic encephalography, cognitive study), cross-sectional population-based study of the mechanisms of cognitive aging spanning adult life expectancy (18-87 years). The dataset includes raw and pre-processed MRI, fMRI, MEG, and cognitive-behavioral data.

### MRI acquisition

All MRI datasets were collected at a single location (MRC-CBSU, Cambridge, England) using a 3T Siemens TIM Trio with a 32-channel head coil. Participants were scanned in a one hour session. The scanner used memory foam cushions for comfort and to minimize head movements.

### MEG acquisition

All MEG data sets were collected at a single location (MRC-CBSU, Cambridge, England) using a 306-channel VectorView MEG system (Elekta Neuromag, Helsinki) consisting of 102 magnetometers and 204 orthogonal planar gradientometers located in a magnetically shielded room (MSR). Data were sampled at 1 kHz with a 0.03 Hz upper-pass filter. Recordings were made in a sitting position. Head position in the MEG helmet was continuously assessed using four head position indicator (HPI) coils (two behind the ears as high as possible without being on the hair, and two on the forehead well separated but not on the hair), to allow autonomous correction of head motion. Two pairs of bipolar electrodes were used to record vertical and horizontal electrooculogram (VEOG, HEOG) signals to monitor blinks and eye movements, and one pair of bipolar electrodes recorded an electrocardiogram (ECG) signal to monitor heart rate-related artifacts. MEG data were collected at rest, participants sat motionless with eyes closed for at least 8 minutes and 40 seconds.

## Data processing

### MEG data

To suppress external noises, we applied Maxwell filter [Taulu and Kajola, 2005; Taulu and Simola, 2006] routine from MNE [Gramfort et al., 2013] package to the raw MEG data. Time series of magnetometer channels were selected for further analysis. On the next step time series were trimmed to 300 seconds length, starting from 100th second of recording, thus producing time series of equal length and in the same frequency grid in all experiments. An offset of 100 seconds was chosen to allow all subjects to enter the resting state. A duration of 300 seconds provides sufficient accuracy for further analysis. Using the FFTW library [Frigo and Johnson, 2005], multichannel spectra were computed over the entire length (300 sec) of the time series. Data structures describing the spatial configuration of the magnetic encephalograph and its position in space relative to the subject’s head were constructed using the FieldTrip [Oostenveld et al., 2011] package.

### MRI data

Usually, individual MRI is used to build a head model for the subsequent solution of the inverse problem of magnetic, limiting the possible solution space to the brain only. In our approach MRI is used for signal/noise separation of the functional tomogram. For this purpose, we extracted the region containing the brain from the MRI using a pre-trained UNesT neural network [Huo et al., 2019; Zhang et al., 2022]. The weights for this network were taken from the Project MONAI [Cardoso et al., 2022] repository. For the purposes of this study, we combined resulting parcellation into a single region of interest, obtaining “brain mask” for every subject’s MRI. Acquired “brain masks” were manually inspected and corrected for errors.

### MEG-based functional tomography

Functional tomography is an MEG analysis technique based on precise frequency decomposition of non-averaged full-length time-series and consequent inverse problem solution without additional constraints. As a result of such analysis, input MEG time-series is converted to a 3-dimensional map of measured activity sources. Each activity source is characterized by its position, direction, measured spectral power and calculated dipole amplitude. Let us consider the theory and experimental justification of the MEG-based functional tomography, as they were set out in the works [Llinás and Ustinin, 2014; Llinás et al., 2015; Ustinin et al., 2023]. The encephalograph simultaneously records the values of magnetic field induction in *K* channels during time *T* producing a set of experimental vectors {*B*_*k*_},*k* = 1,…,*K*. The field is recorded at *H* points in time with a constant step *T/H*. As a result of the measurements, we obtain a two-dimensional data array *B*_*hk*_, where *h* = 0,…,*H* −1,*k* = 1,…,*K*. Multichannel discrete Fourier transform can be written as:

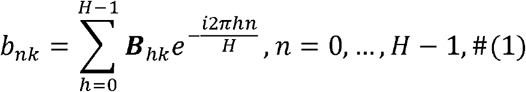

where *b*_*nk*_ – complex Fourier amplitude for frequency *v*_*n*_ in a channel with a number *k*, and 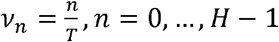. All spectra are computed for the full measurement time *T*, which reveals the detailed frequency structure of the system. Frequency resolution is equal to *δv* = *v*_*n*_ − *v*_*n*-1_ = 1/*T*, thus, the frequency resolution is determined by the registration time. The phase and amplitude of the *n*-th component of the Fourier series are written as: 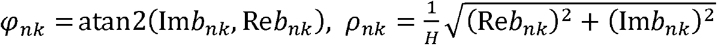, where *Reb*_*nk*_ – real part of *b*_*nk*_, Im*b*_*nk*_ – imaginary part.Knowing the multichannel spectrum, the inverse Fourier transform can be performed on all channels:

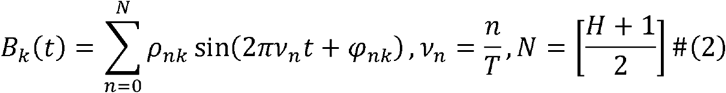

In order to study the detailed frequency structure of the brain, we reconstruct the multi-channel signal at each frequency and analyze the obtained time series. Reconstructed multichannel signal of frequency *v*_*n*_ in all channels:

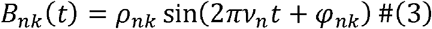

where 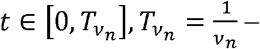 period of that frequency.

To describe the system under study, we adopt a model in which one frequency component is described by one current dipole. We’ve found, that the magnitude ratios between channels remain constant, only the scale changes. Thus formula (3) can be written as:

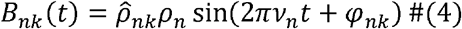

where 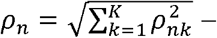 amplitude, 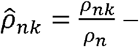 value of normalized pattern of this oscillation in *k*-th channel. In multichannel measurements, the space is determined by the channel arrangement. The use of normalized patterns makes it possible to determine the spatial structure of the signal sources by the solution of the inverse problem, and this structure remains constant over the entire oscillation time. The time dependence of the field is determined by the function *ρ*_*n*_ sin(2*πv*_*n*_*t* + Φ_*n*_), which is common for all channels, i.e., this source oscillates as a single unit at frequency *v*_*n*_.

The theoretical foundations for the reconstruction of static functional entities (neural circuits, or sources) were outlined in [Llinás and Ustinin, 2014; Llinás et al., 2015]. This reconstruction is based on detailed frequency analysis and the selection of frequency components with similar patterns. Algorithm for this method can be written as:

1. Fourier transform of the input multichannel signal.
2. Inverse Fourier transform – reconstruction of the signal at each frequency.
3. Calculation of phase, normalized pattern and its amplitude for each frequency component.
4. Solution of the inverse problem for all of the components.

To calculate phases, normalized patterns of experimental signal components and their amplitudes we use the following procedure for each frequency:

1. Reconstruct a multichannel signal at a single frequency for a single period of that frequency.
2. Find a maximum of that reconstructed signal. For this purpose, we calculate the sum of squares of amplitudes in all channels at each point in time.
3. At the point of maximum, we take a “slice” of the multichannel signal, representing a vector of length k, and calculate the norm of this vector. This normalized vector is a pattern 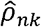 of selected frequency.
4. Phase estimation for this frequency:

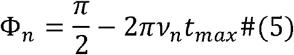

where *t*_*max*_ is moment of maximum, found on step 2.

Each elementary oscillation is characterized by frequency *v*_*n*_, phase Φ_*n*_, amplitude *ρ*_*n*_, normalized pattern 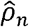, and its source is a functional entity with a permanent spatial structure. The functional tomography method reconstructs the structure of the system by analyzing a set of normalized patterns 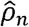. A functional tomogram shows a three-dimensional map of the distribution of energies produced by sources located at a given position in space. To construct a functional tomogram, the investigated region of space is divided into *N*_*x*_ × *N*_*y*_ × *N*_*z*_ elementary cubic cells with centers at *r*_*ijs*_. The length of the cube edge is chosen according to the desired accuracy, in this study it was 2 mm. In each cell set of test dipoles *Q*_*ijsl*_, |*Q*_*ijsl*_ |= 1*nAm* with direction ***l*** is constructed. The magnetic induction generated by the test dipole *Q*_*ijsl*_, located at the point *r*_*ijs*_ is recorded by a sensor with the number *k*, located at the point with coordinates *r*_*k*_ and having the direction *n*_*k*_, *k*-th component 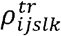 of the test pattern 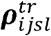 is calculated using the model of the current dipole in a spherical conductor [Sarvas, 1987]:

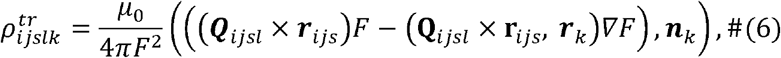

Where 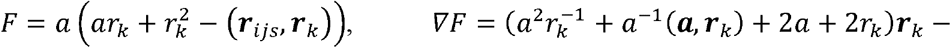 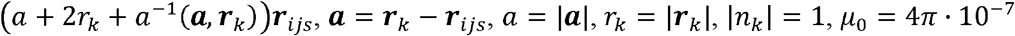.

The normalized pattern is calculated as:

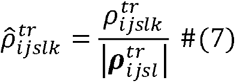

where 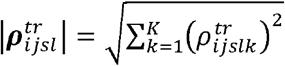. All test dipoles located at the point *r*_*ijs*_, lie within the same plane orthogonal to *r*_*ijs*_, since the result of the vector product *Q*_*ijsl*_ × *r*_*ijs*_ is non-zero only for such dipoles. The test dipoles cover the circle in *L*_*max*_ directions in 360/*L*_*max*_ degree increments, *L*_*max*_ = 72 was used in this study. For each of the dipoles, a set of normalized patterns is computed using formula (7):

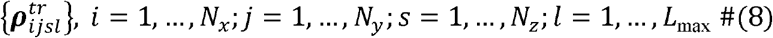

For each of the normalized experimental patterns 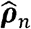 the following function is computed to determine the difference between that pattern and one of the test patterns:

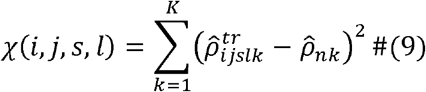

where 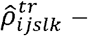 *k*-th component of the test pattern *ijsl*, 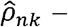 *k*-th component of the normalized experimental pattern *n, k* – channel number. Position and direction of the source corresponding to the pattern 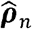, is determined by the numbers (*I, J, S, L*), corresponding to the minimum of the function χ(*i,j,s,l*) on variables *i* = 1,…, *N*_*x*_; *j* = 1,…, *N*_*y*_; *s*= 1,…, *N*_*z*_; *l* = 1,…, *L*_max_. The minimum of this function is found by the method of exhaustive search, i.e., by selecting the smallest of several million values of the function χ for each pattern 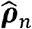. This procedure determines the position *r*_*IJS*_ – solution of the inverse problem for pattern 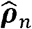, without spatial filtering of channels and without introduction of weight functions. In this study, we propose a method for direct estimation of the amplitudes of the dipole sources producing the measured magnetic field:

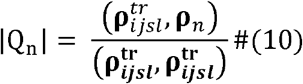

Repeating this procedure for all normalized patterns 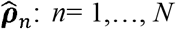, it is possible to find position, direction and amplitude of the source dipole for all frequency components of the initial experimental signal, thus solving the inverse problem of magnetic encephalography. After solution of the inverse problem, a functional tomogram, a three-dimensional distribution of sources in space, is obtained. Each cell of space corresponds to a set of sources characterized by their frequencies, dipole moments and their spectral powers.

The electric power of the elementary dipolar source can be defined as total energy, produced by this source per unit time:

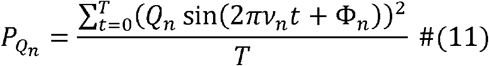

Electric power of the brain in particular frequency band is a sum of powers of all sources located in the brain, whose frequency lies in this band.

### Data processing and analysis

Functional tomograms were computed for all 595 subjects’ recordings with the following parameters: time series duration 300 sec, frequency bandwidth 1−100Hz, spatial resolution 2mm, number of directions 72. On the next step of analysis each functional tomogram was separated into two parts: one containing sources localized in the brain, other containing all other physiological sources. Such separation was achieved by combining functional tomogram and whole brain mask, acquired on MRI data preprocessing stage [Llinás et al., 2022]. After that time series were reconstructed for all brain functional tomograms and total electric energies were calculated. The outliers were found by median filtering. The energies of these outliers differed from the median values by several orders of magnitude. These experimental data were discarded from further analysis. This procedure left 501 experimental sets. Then remaining experimental sets were divided by participants sex and arranged in 10 years cohorts. Numbers of participants in each resulting cohorts were equalized, resulting in selection of 434 sets: 217 for males and 217 for females. Numbers of participants in each cohort and their mean age are presented in Table 1.

**Table 1.**
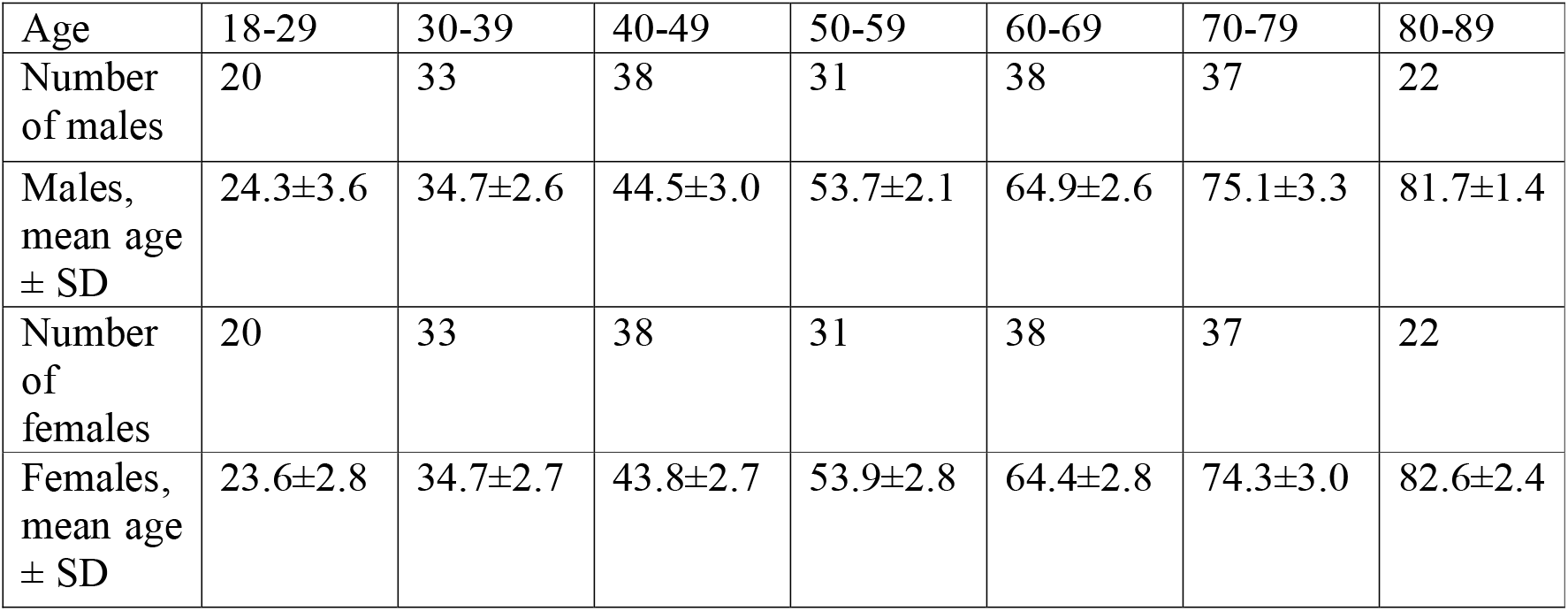
Distribution of number of experiments by age cohorts and sex.

## Results

Let us consider results, obtained for sex- and age-related differences in the electric power of the brain. Figure 1 presents an overview of the age dependencies found for females’ absolute average power (left) and males’ absolute average power (right). Linear regression coefficients are shown in Table 2.

**FIGURE 1.**
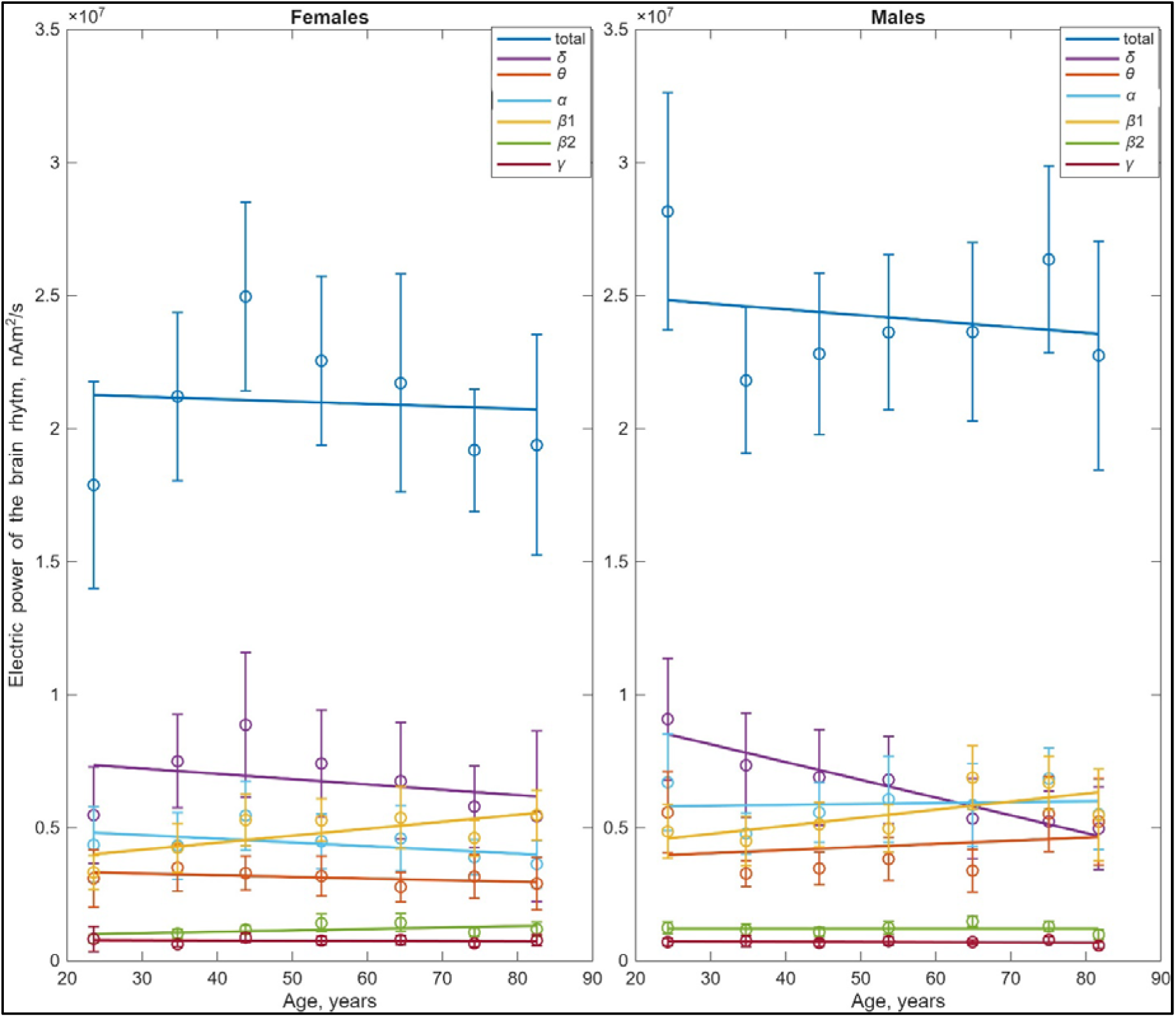
Dependencies of average electric power on age for females (left) and males (right), all brain rhythms. Colors corresponding to rhythms are indicated at the subpanels.

**Table 2.** Linear regression.

|  |  | Rhythm |  |  |  |  |  |
| --- | --- | --- | --- | --- | --- | --- | --- |
|  |  | Delta | Theta | Alpha | Beta1 | Beta2 | Gamma |
| Males | a0 | 9.69E+06 | 3.51E+06 | 5.81E+06 | 3.63E+06 | 1.10E+06 | 7.31E+05 |
|  | a1 | -5.91E+04 | 1.43E+04 | 1.26E+03 | 3.49E+04 | 2.22E+03 | -7.09E+02 |
|  | p-val | **0.02** | 0.59 | 0.93 | **0.01** | 0.28 | 0.56 |
| Females | a0 | 6.96E+06 | 3.59E+06 | 5.41E+06 | 3.48E+06 | 8.58E+05 | 7.26E+05 |
|  | a1 | -2.30E+03 | -7.97E+03 | -1.84E+04 | 2.42E+04 | 5.38E+03 | 1.91E+02 |
|  | p-val | 0.93 | 0.09 | 0.08 | 0.06 | 0.23 | 0.90 |

As follows from Figure 1, total brain power for females is lower than such for males and demonstrates a different age trajectory. Rhythms corresponding to lower frequencies from delta to beta 1 (1-21 Hz) contribute similar amounts, between 20 and 30 percent each. The two high-frequency rhythms, beta 2 and gamma (21-48 Hz), together account for 8-9 percent of the total power.

Absolute values of the total electric power are presented in Figure 2. The drastic difference can be seen in the youngest cohort (18-29 years), females’ power is 65% of such for males. In the next adult cohort (30-39 years), in the middle cohorts, (40-49 years and 50-59 years) and in the first old cohort (60-69 years) males’ and females’ powers are approximately equal (93% to 110%). For two oldest cohorts the females’ power is lower than males’ (74% for 70-79 years and 85% for 80-89 years).

**FIGURE 2.**
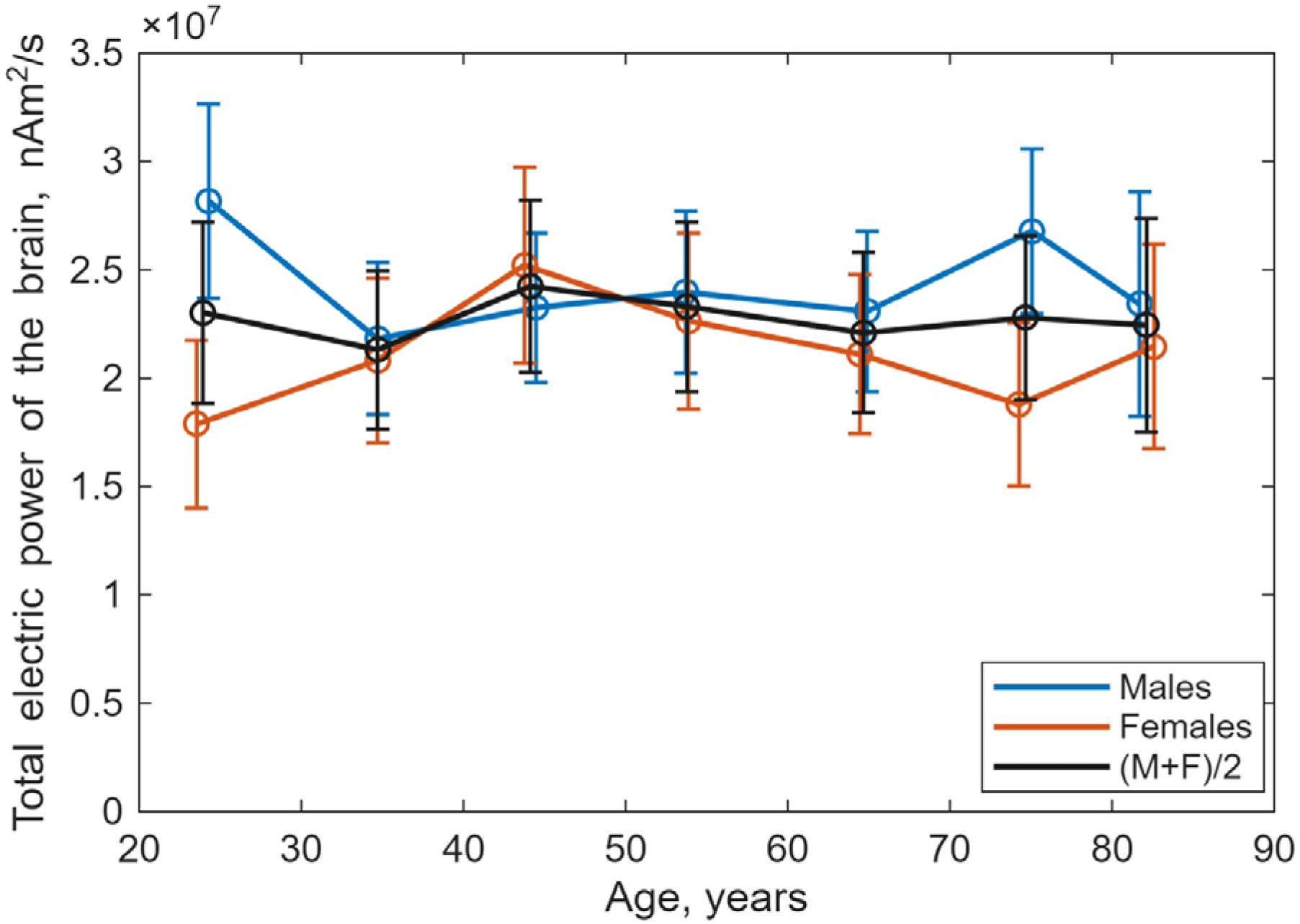
Dependencies of average electrical power on age for females (red) and males (blue), compared with intersex average (black), for total brain power (1-48 Hz).

If we assume that electrical sources in the brain are, on average, distributed uniformly, then their total power will be approximately proportional to the radiating volume. Average total power through the lifespan for males is 116.4% of such for females. This result is in reasonable agreement with the anatomical data from [Gur, 1999], where the sum of gray matter and white matter volumes for males was found to be 114.7% of that for females.

The Figure 2 also presents the result of the intersex averaging (black color), which is the simple sum of the male and female curves, divided by 2. This operation can be performed correctly, because the number of males is equal to the number of females in each cohort. One can see, that such average curve is much smoother than each of males’ and females’, and represents almost constant electric power through the lifespan. This is in good agreement with the results of the article [Ustinin et al, 2025], where it was found that the electric power of the mixed F+M population is constant throughout a person’s life. Note that in the article [Ustinin et al, 2025], the ratio of males to females in each age cohort was a random variable ranging from 0.55 to 1.37. However, some averaging did occur in that study as well. To investigate this effect, in the current study, we included an equal number of males and females in each cohort.

Considering the known facts about the anatomical differences between the brains of males and females [Lenroot, 2010, Ritchie, 2018, Ruigrok, 2014, Bethlehem, 2022, Vucic, 2026, Ballard, 2022, Gur, 1999], it seems reasonable to study the sex-related and age-related characteristics of the relative shares of various rhythms in total electric power. Let us compare the age-related percentage shares of various rhythms in the total electrical power of the brain for females and males, see Figure 2 and Tables 3-4.

**Table 3.** Females’ percentage of the brain rhythms electric power in the age cohorts.

|  |  | Age cohort |  |  |  |  |  |  |
| --- | --- | --- | --- | --- | --- | --- | --- | --- |
|  |  | 18-29 | 30-39 | 40-49 | 50-59 | 60-69 | 70-79 | 80-89 |
| % of electric power in band | **Delta** | 30.6 | 34.2 | 36.2 | 33.2 | 29.1 | 28.7 | 34.9 |
|  | **Theta** | 17.3 | 16.8 | 13.1 | 14.1 | 13.1 | 16.9 | 13.5 |
|  | **Alpha** | 24.4 | 20.8 | 21.6 | 19.8 | 21.8 | 20.7 | 16.9 |
|  | **Beta1** | 18.6 | 20.4 | 21.0 | 23.4 | 25.5 | 24.5 | 25.5 |
|  | **Beta2** | 4.6 | 4.9 | 4.7 | 6.3 | 6.8 | 5.7 | 5.5 |
|  | **Gamma** | 4.5 | 3.0 | 3.4 | 3.4 | 3.7 | 3.5 | 3.6 |

**Table 4.** Males’ percentage of the brain rhythms electric power in the age cohorts.

|  |  | Age cohort |  |  |  |  |  |  |
| --- | --- | --- | --- | --- | --- | --- | --- | --- |
|  |  | 18-29 | 30-39 | 40-49 | 50-59 | 60-69 | 70-79 | 80-89 |
| % of electric power in band | **Delta** | 32.2 | 33.7 | 31.6 | 29.8 | 20.7 | 21.2 | 24.1 |
|  | **Theta** | 19.8 | 15.0 | 14.9 | 15.9 | 14.7 | 20.6 | 22.3 |
|  | **Alpha** | 23.8 | 21.9 | 23.9 | 25.3 | 25.4 | 25.6 | 23.5 |
|  | **Beta1** | 17.3 | 20.6 | 22.0 | 20.7 | 29.8 | 24.9 | 23.4 |
|  | **Beta2** | 4.4 | 5.3 | 4.7 | 5.2 | 6.4 | 4.8 | 4.2 |
|  | **Gamma** | 2.5 | 3.4 | 2.9 | 3.1 | 3.0 | 2.9 | 2.5 |

As follows from Figure 3 and Tables 3-4, general age dependences of relative shares are qualitatively similar for males and females, while some sex-related differences may be noted.

**FIGURE 3.**
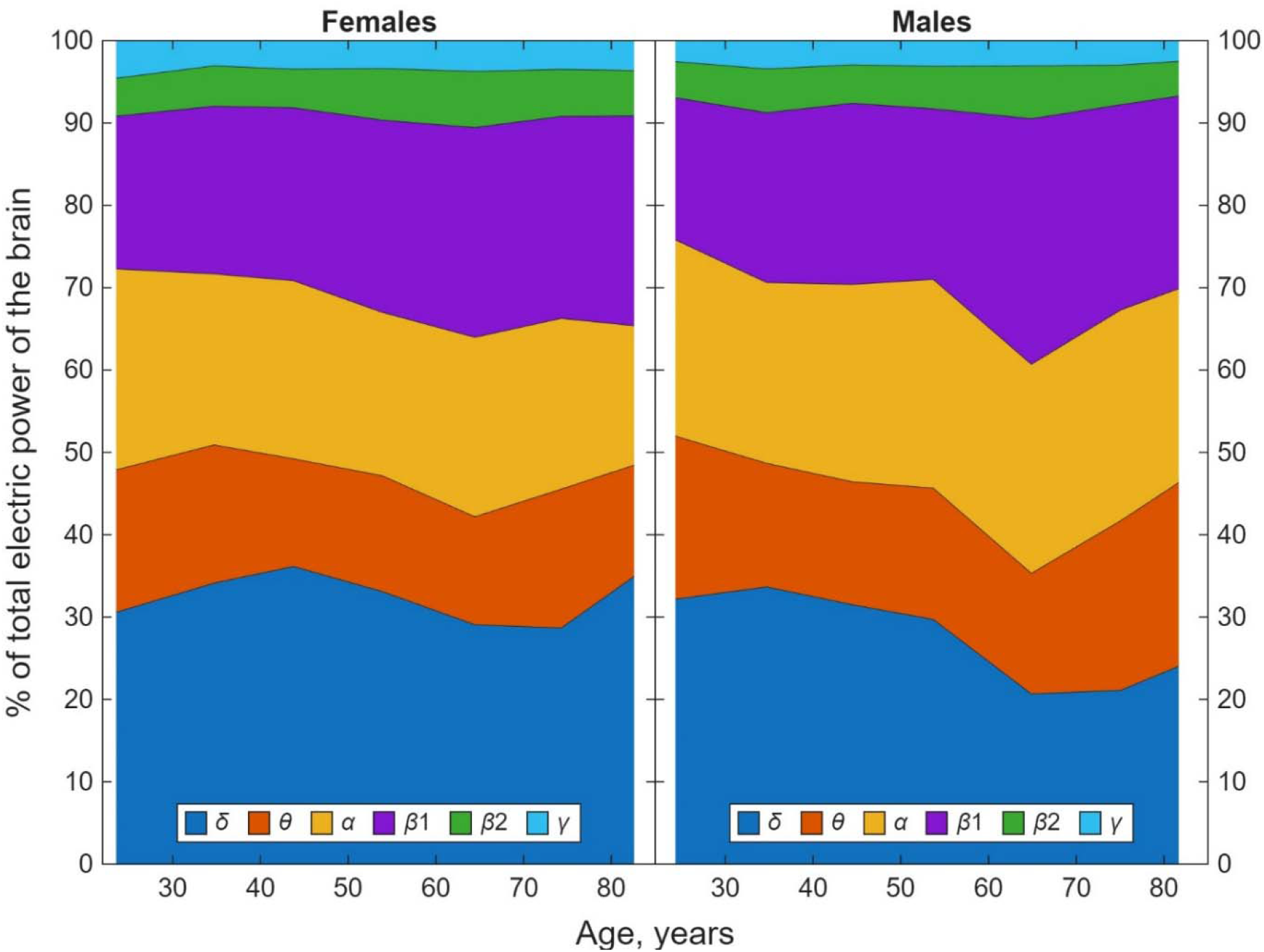
Percentage of power of different brain rhythms depending on age for females (left) and males (right), all brain rhythms. Colors corresponding to rhythms are indicated at the subpanels.

### 1. Delta rhythm (1-4 Hz)

**Absolute** values of the delta power are presented in Figure 4A. The drastic difference can be seen in the youngest cohort (18-29 years), females’ power is 60% of such for males. In the next adult cohort (30-39 years) the females’ power is 2% higher, and in the first middle cohort (40-49 years) it is 28% higher. Starting from the second middle cohort (50-59 years) up to the oldest cohorts the females’ power is always higher, from 24% to 10%. Average delta power through the lifespan for males is 94.7% of such for females.

**FIGURE 4.**
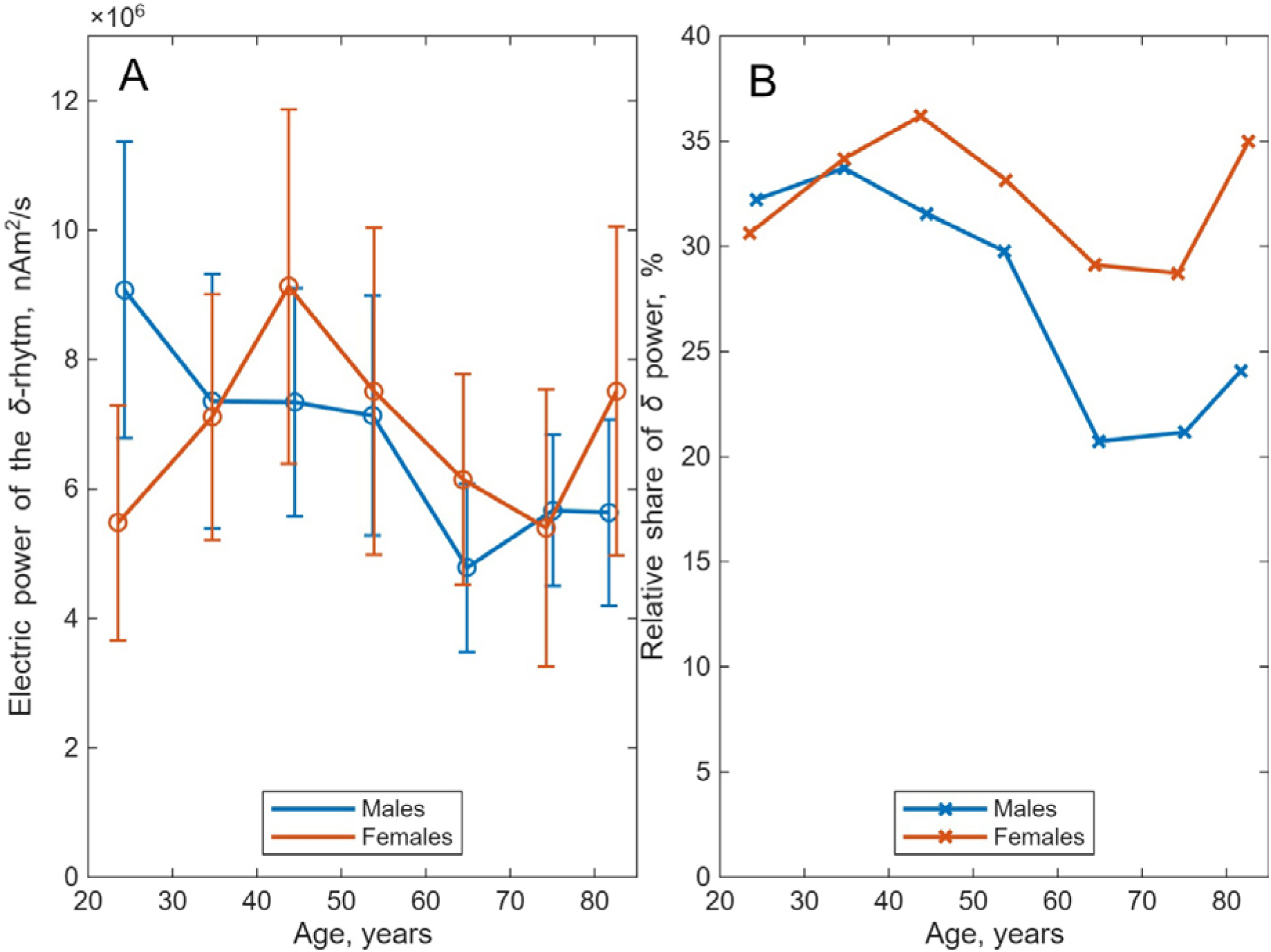
Dependencies of average electrical power on age for females (red) and males (blue), delta rhythm power (1-4 Hz), Panel A - absolute values, panel B – relative shares of total power.

**Relative** shares of the delta power are presented in Figure 4B. The percentages of the delta rhythm power are approximately equal, δ_f_/δ_m_=0.98, for males and females in younger cohorts (18-29 years and 30-39 years). In the middle age (40-49 years and 50-59 years) the female’s percentage of delta power is somewhat higher, δ_f_/δ_m_=1.13, and in the oldest cohorts (60-69 years, 70-79 years and 80-89 years) the percentage for females significantly exceeds such for males: δ_f_/δ_m_=1.4. The delta power percentage for females is oscillating from 29% to 36% through the lifespan, being almost maximal (35%) in the oldest cohort (80-89 years), with the lifetime average 32.4%. For males the delta power percentage decreases with age from 34% to 24% with minimum in the oldest cohort, with the lifetime average 27.6%.

Similar results are described in [Woodhouse, 2025], obtained by the intracranial EEG. In this paper one can see the linearly falling delta rhythm, equal for males and females, while in our work delta rhythm for females prevails over such for males in all cohorts, except first and second.

### 2. Theta (4-8 Hz)

**Absolute** values of the theta power are presented in Figure 5A. The drastic difference can be seen in the youngest cohort (18-29 years), females’ power is 54% of such for males. In the next adult cohort (30-39 years) the females’ power is 10% higher. Starting from the first middle cohort (40-49 years) to the first older cohort the females’ power slowly falls from 93% to 81% of males’, and in the two oldest cohorts it is again rather small, 57%. Average theta power through the lifespan for males is 133.4% of such for females.

**FIGURE 5.**
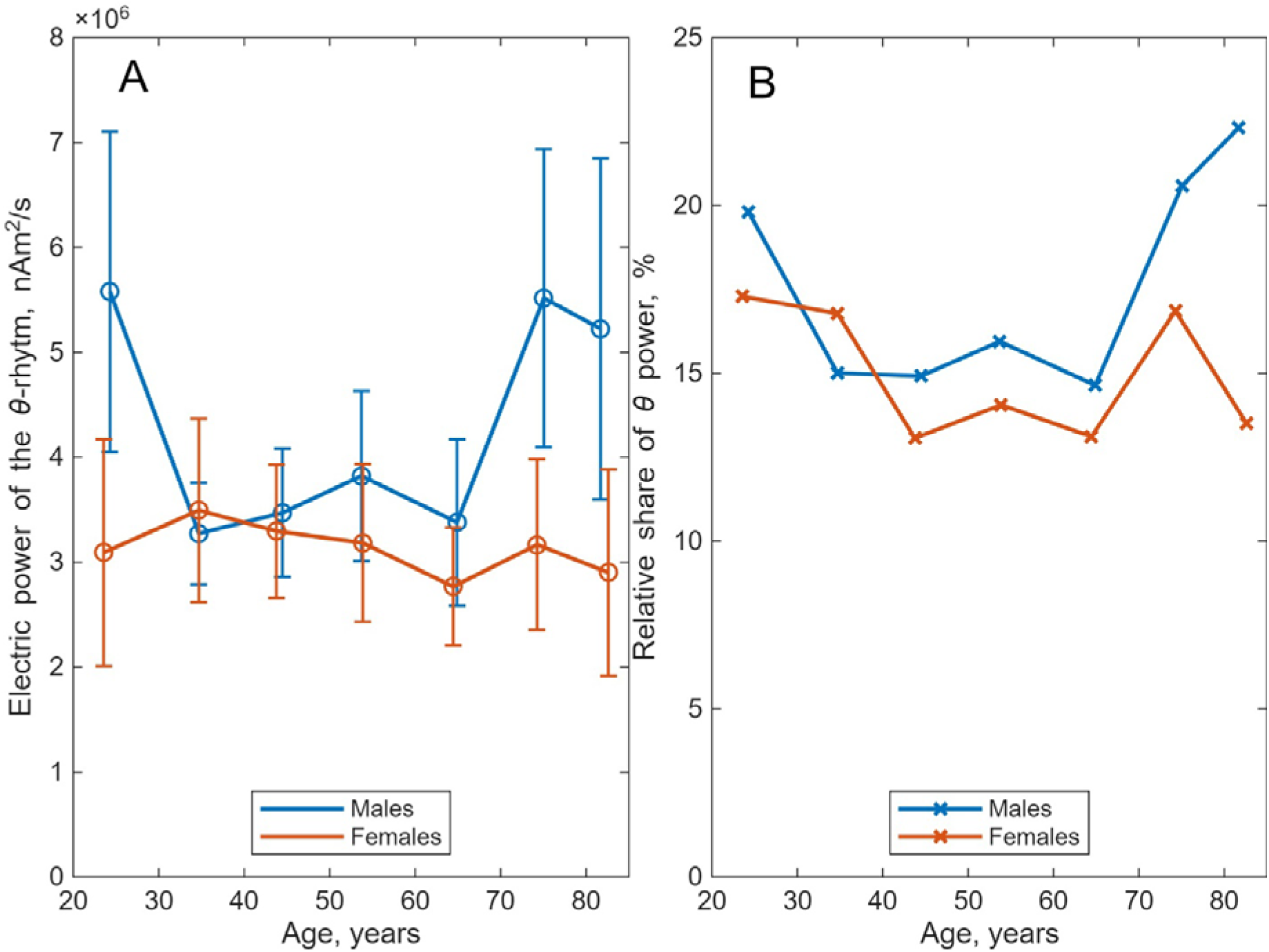
Dependencies of average electrical power on age for females (red) and males (blue), theta rhythm power (4-8 Hz). Panel A - absolute values, panel B – relative shares of total power.

**Relative** shares of theta power are presented in Figure 5B. The percentage of the theta rhythm power for females is lower in the youngest cohort (18-29 years), θ_f_/θ_m_ =0.87. In the second cohort (30-39 years) the females’ percentage is higher, θ_f_/θ_m_ =1.12. In cohorts from 40 to 79 years the females’ percentage is lower, θ_f_/θ_m_ =0.87, while in the oldest cohort (80-89 years) it is minimal θ_f_/θ_m_ =61. The theta power percentage for females is oscillating from 17% to 13% through the lifespan, with the lifetime average 15%. For males the theta power percentage oscillates from 15% to 22% through the lifespan, with the lifetime average 17.6%.

Intracranial EEG [Woodhouse, 2025] for theta are falling slower than delta, still equal for males and females, while in our work theta rhythm for males prevails over such for females.

### 3. Alpha (8-13 Hz)

**Absolute** values of the alpha power are presented in Figure 6A. The drastic difference can be seen in the youngest cohort (18-29 years), females’ power is 65% of such for males’. In the two next adult cohort (30-49 years) the females’ power is 87% of such for males’ and in the four older cohorts (50-89 years) the females’ power is 69% of such for males’. Average alpha power through the lifespan for males is 132.1% of such for females.

**FIGURE 6.**
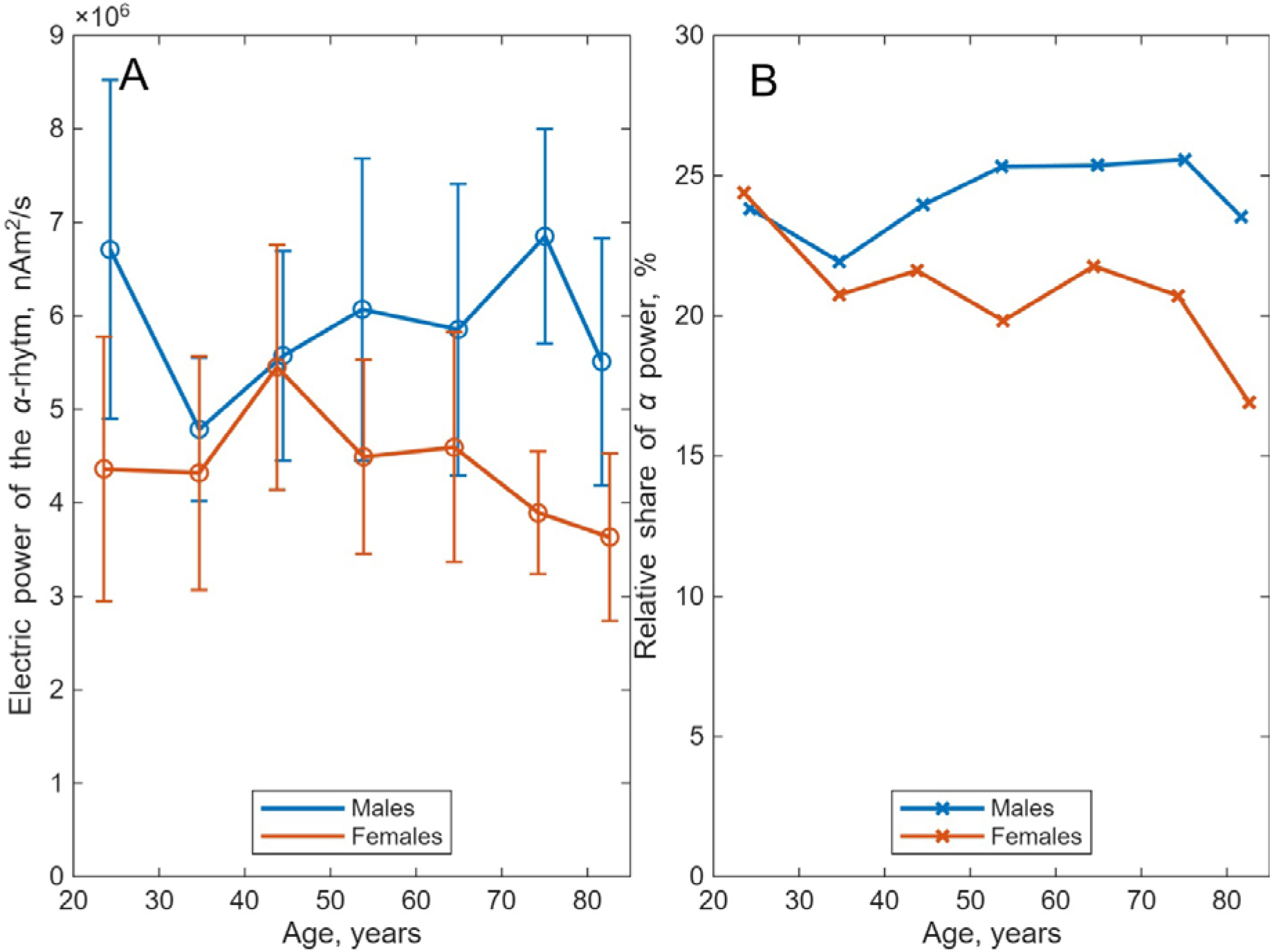
Dependencies of average electrical power on age for females (red) and males (blue), alpha rhythm power (8-13 Hz). Panel A - absolute values, panel B – relative shares of total power.

**Relative** shares of alpha power are presented in Figure 6B. The percentages of the alpha rhythm power are approximately equal, α_f_/α_m_=0.95, for males and females in three younger cohorts (from 18 to 49 years). In four older cohorts (from 50 to 89 years) the female’s percentage of alpha power is lower, α_f_/α_m_=0.8. The alpha power percentage for females is falling from 24.4% to 16.9% through the lifespan, with the lifetime average 20.9%. For males the alpha power percentage oscillates with age from 21.9% to 25.6% with the lifetime average 24.2%.

Intracranial EEG [Woodhouse, 2025] for alpha are slowly growing, identically for males and females, while in our work alpha rhythm for males substantially prevails over such for females.

### 4. Beta1 (13-21 Hz)

**Absolute** values of the beta1 power are presented in Figure 7A. The biggest difference can be seen in the youngest cohort (18-29 years), females’ power is 68% of such for males’. In the three next adult cohorts (30-59 years) the females’ power is equal to such for males’. In the two next cohorts (60-79 years) the females’ power is 74% of such for males’, and in the oldest cohort (80-89 years) the females’ power is equal to such for males’. Average beta1 power through the lifespan for males is 114.6% of such for females.

**FIGURE 7.**
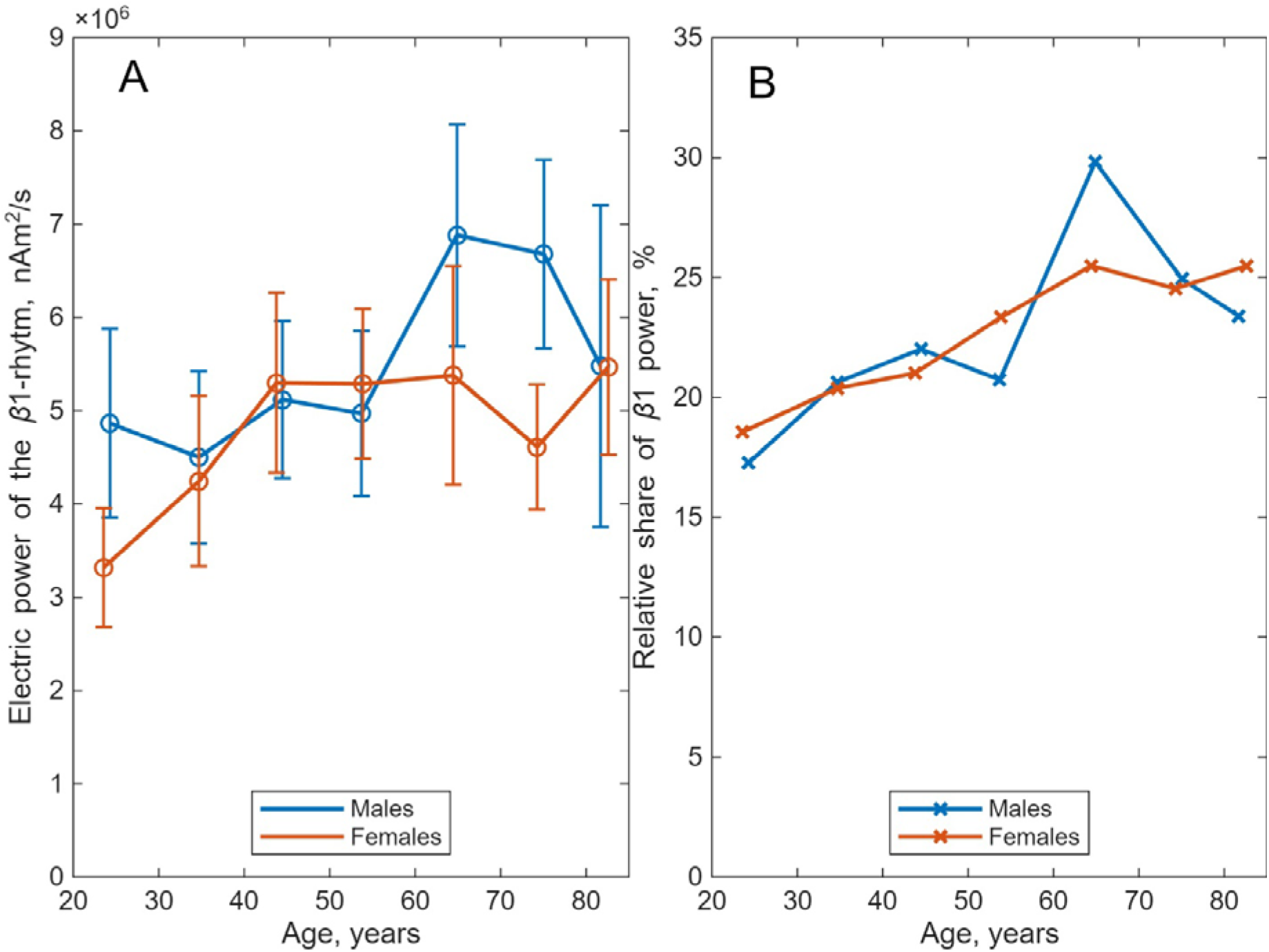
Dependencies of average electrical power on age for females (red) and males (blue), beta1 rhythm power (13-21 Hz). Panel A - absolute values, panel B – relative shares of total power.

**Relative** shares of beta1 power are presented in Figure 7B. The percentages of the beta1 rhythm power are almost equal, β1_f_/β1_m_=1.01, for males and females in all cohorts (from 18 to 89 years). The beta1 power percentage for females is growing from 18.6% to 25.5% through the lifespan, with the lifetime average 22.7%. For males the beta1 power percentage grows with age from 17.3% to 25.6% with the lifetime average also 22.7%.

Intracranial EEG [Woodhouse, 2025] for beta are also slowly growing, identically for males and females, similarly with our results.

### 5. Beta2 (21-30 Hz)

**Absolute** values of the beta2 power are presented in Figure 8A. The biggest difference can be seen in the youngest cohort (18-29 years), females’ power is 67% of such for males’. In the five next cohorts (30-79 years) the females’ power is oscillating around such for males’ with average 98%. In the oldest cohort (80-89 years) the females’ power is 119% of such for males’. Average beta2 power through the lifespan for males is 103.4% of such for females.

**FIGURE 8.**
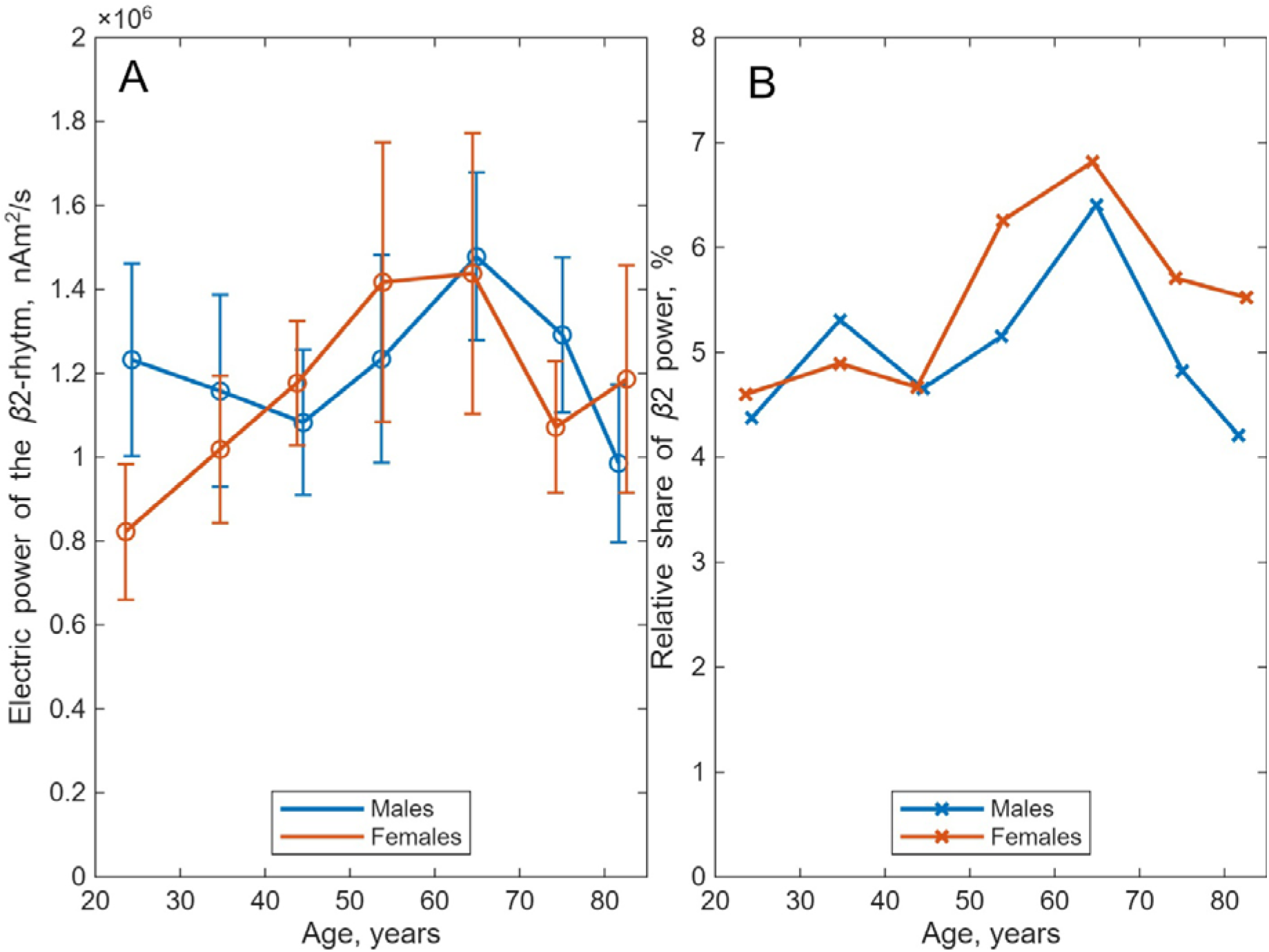
Dependencies of average electrical power on age for females (red) and males (blue), beta2 rhythm power (21-30 Hz). Panel A - absolute values, panel B – relative shares of total power.

**Relative** shares of beta2 power are presented in Figure 8B. The percentages of the beta2 rhythm power are almost equal, β2_f_/β2_m_ =0.99, for males and females in first three cohorts (from 18 to 49 years). In four older cohorts (from 49 to 89 years), the beta2 power percentage for females is higher than such for males’, β2_f_/β2_m_ =1.19. The beta2 power percentage for females oscillates from 4.6% to 6.8% through the lifespan, with the lifetime average 5.5%. For males the beta2 power percentage oscillates from 4.2% to 6.4% with the lifetime average 5.0%.

The article [Woodhouse, 2025] considers beta as a wide range 13-30 Hz, so beta 2 cannot be directly compared.

### 6. Gamma (30-48 Hz)

**Absolute** values of the gamma power are presented in Figure 9A. Age dependencies oscillate around each other, with a maximal difference in the oldest cohort (80-89 years), where the females’ power is 141% of such for males’. Average gamma power through the lifespan for males is 94.9% of such for females.

**FIGURE 9.**
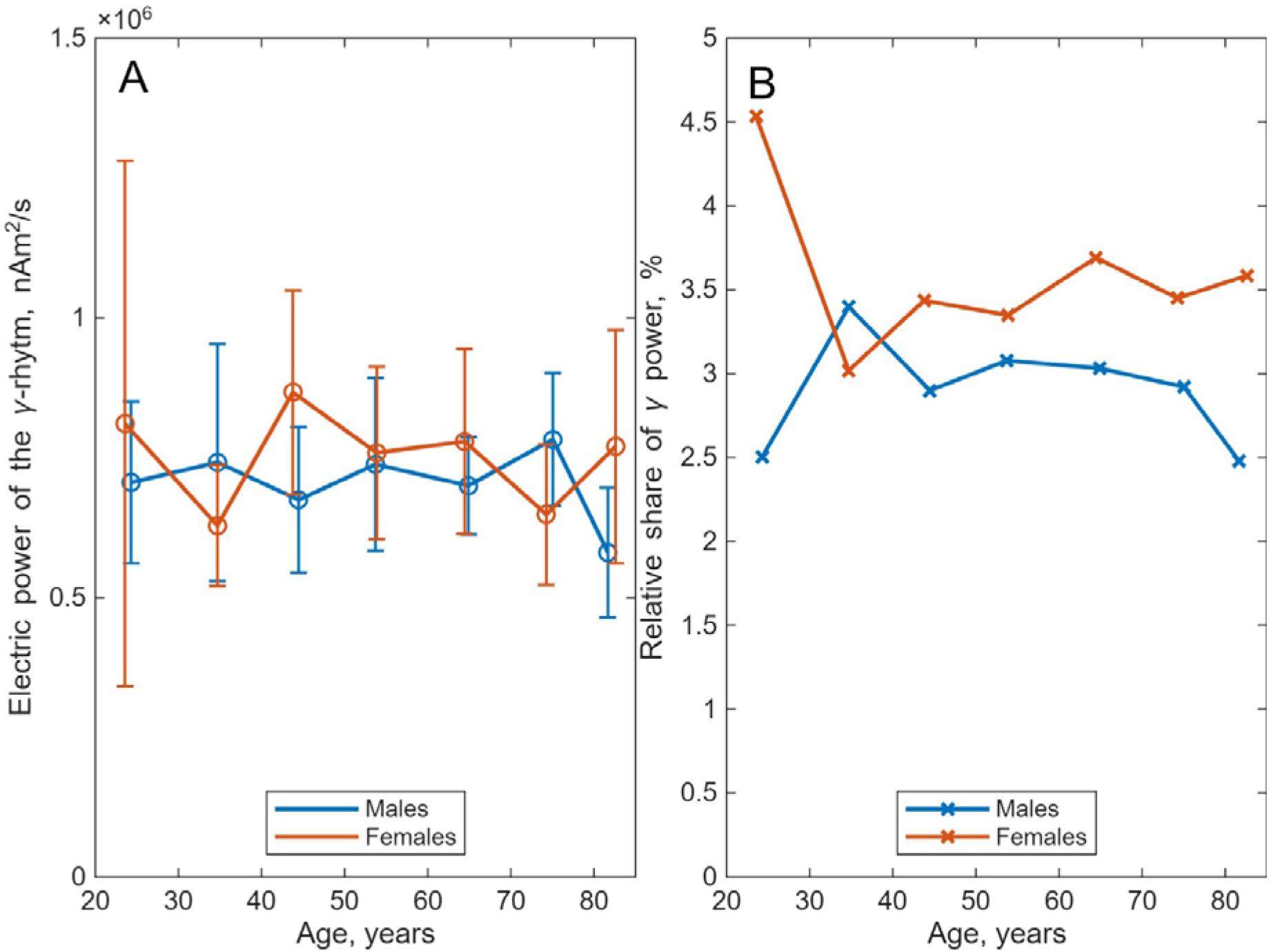
Dependencies of average electrical power on age for females (red) and males (blue), gamma rhythm power (30-48 Hz). Panel A - absolute values, panel B – relative shares of total power.

**Relative** shares of gamma power are presented in Figure 9B. The percentage of the gamma rhythm females’ power is generally higher than such for males’, average γ_f_/γ_m_ =1.12 for five middle cohorts (from 30 to 79 years). There are drastic differences in the first cohort (18 to 29 years), γ_f_/γ_m_ =1.80, and in the last cohort (80 to 89 years) γ_f_/γ_m_ =1.44. The gamma power percentage for females oscillates from 3.0% to 4.5% through the lifespan, with the lifetime average 3.6%. For males the gamma power percentage oscillates from 2.5% to 3.4% with the lifetime average 3.0%.

Similar results are described in [Woodhouse, 2025], obtained by the intracranial EEG for the band (30-77.5 Hz). In this paper one can see the constant gamma rhythm, with prevailing females’ curve. We obtained the same results, while considering more narrow range for gamma (30-48 Hz).

## CONCLUSION

The work examined the sex and age dependencies of the electrical power of the human brain. Age-related trajectories of relative rhythm powers (normalized to total brain power) were found to show qualitative agreement between males and females (see Figure 3). A key advantage of our proposed method is that it allows for direct measurement of the electrical power of elementary sources in the brain. This opens up the possibility of using the method as a data source for normative studies of brain function, diagnostics, and drug evaluation. Therefore, it is important to study the absolute values of electrical power and its dependence on age and sex. We found that age-related changes in absolute values of electrical power differ significantly between males and females in four of the six main brain rhythms: delta, theta, alpha, and beta 1. These rhythms correspond to frequencies from 1 to 21 Hz and contain over 90% of the total brain power in both males and females (See Figures 4-7). If we’re studying phenomena affecting these rhythms, mixed-sex samples are meaningless.

Also, from the results of the comparison of the age dependencies of various rhythms of males and females and from Figure 2, an important methodical question follows about the constructing correct sample populations. Specifically, the summary curve in Figure 2 (black) is unrelated to either males or females. It describes the average lifetime trajectory of total electrical brain power for a sample with equal numbers of males and females. When using brain electrical power as a normative parameter, it is necessary to form samples that are homogeneous by sex.

## Data and code availability

Raw data are available at https://camcan-archive.mrc-cbu.cam.ac.uk/dataaccess/ under specified conditions. Code used for the pre-processing and analyses is available at https://osf.io/2r6fb/.

## Funding

This work was supported by Moscow Center of Fundamental and Applied Mathematics, Agreement with the Ministry of Science and Higher Education of the Russian Federation, No. 075-15-2025-346.

## Conflict of interest

The authors declare that they have no conflict of interest.

## Notes

https://osf.io/2r6fb/

https://camcan-archive.mrc-cbu.cam.ac.uk/dataaccess/

### Competing Interest Statement

The authors have declared no competing interest.

